# Reach corrections toward moving objects are faster than reach corrections toward jumping targets

**DOI:** 10.1101/2022.12.24.521682

**Authors:** Sasha Reschechtko, Cynthiya Gnanaseelan, J. Andrew Pruszynski

## Abstract

Visually guided reaching is a common motor behavior that engages subcortical circuits to mediate rapid corrections. These circuits help us successfully grasp objects we see, even if those objects move during the reach. Although these neural mechanisms have evolved for interaction with the physical world, they are almost always studied in the context of reaching toward virtual targets displayed on a computer or projection screen. Virtual targets are unrealistic both because they cannot be physically grasped and because they generally move by “jumping” from one place to another instantaneously. Recent work has indicated that various aspects of “real” visual stimuli, including proximity and graspability, elicit distinct neural activity. In this study, we instructed participants to perform rapid reaches to physical objects. On some trials, these objects either moved realistically (continuously from one position to another) or unrealistically (jumping instantaneously to an equivalent position). Participants were consistently faster in correcting their reach trajectories when the object moved continuously.

## Introduction

Visually guided reaching is an extremely common activity in everyday life. However, in many cases we reach toward objects that are not stationary -- like when we reach to pet a cat, only for it to quickly move to avoid our affection. Many groups have studied how quickly we can react to sudden changes in a reach target, finding that we are able to change an ongoing reaching movement quite quickly, with kinematic changes occurring less than 150 ms following the change of target position (Brenner & Smeets, 1997; Day & Lyon, 2000; Franklin & Wolpert, 2008; Gu et al., 2016; Paulignan et al., 1991; Pélisson et al., 1986; Prablanc & Martin, 1992; Pruszynski et al., 2010). These corrections do not appear to rely on conscious perception of hand position (Goodale et al., 1986) involve subcortical pathways (Alstermark et al., 1987; Day & Brown, 2001). These corrections are often considered automatic because they guide the hand to a moving target even if this conflicts with verbal instructions to do something else when the target moves, e.g. move in the other direction (Day & Lyon, 2000) or stop an ongoing movement (Pisella et al., 2000).

Although the motivation for reach-correction studies often relates to the ability of the motor system to intercept physical objects, very few studies have investigated reach corrections with actual physical objects (Battaglia-Mayer et al., 2013; Blouin et al., 1993; Camponogara & Volcic, 2019; D’Mello et al., 1985; Shaw et al., 2022) Instead of reaching toward physical objects, participants in most studies reach toward images on a computer or projection screen (“virtual targets”). Although physical objects are less convenient to use in laboratory settings than virtual targets, the motor system is likely specialized to interact with physical objects. In fact, a growing body of literature indicates that interacting with virtual representations of objects is often not a good proxy for interaction with the physical objects themselves (Snow & Culham, 2021). For example, neural activity is modulated according to whether or not objects are physical (Freud et al., 2018) and even whether or not a physical object is close enough to the body for interaction (Fairchild et al., 2021; Gallivan et al., 2009). Additionally, when planning interactions with real objects compared to virtual targets or unreachable objects, the surrounding environment is taken into account to a greater extent, potentially indicating that physical interaction is planned to a greater extent, if the object is actually reachable (Gomez et al., 2018).

We recently described aspects of rapid reach corrections under tactile guidance using an oriented edge or the relative motion of a textured surface under the fingertip (Pruszynski et al., 2016; Reschechtko & Pruszynski, 2020). These studies involved reaching toward a physical object in order to physically link it to the tactile stimulus. This physical object paradigm differs from most previous reaching paradigms for two main reasons. First, as previously mentioned, the target of the reach is a physical object; second, as a physical object, the target moves in a physically possible way, i.e. continuously, rather than disappearing in one location and instantaneously reappearing in another location (“jumping”).

Using virtual targets does not prevent those targets from moving in a continuous manner. Nonetheless, most paradigms for reaching updates involve instantaneous target movement (Day & Lyon, 2000; Prablanc & Martin, 1992; Smeets & Brenner, 2003; Soechting & Lacquaniti, 1983), instantaneous movement of a background to affect the visual impression of target movement (Gomi et al., 2006; Saijo, 2005), or instantaneous movement of the cursor representing the hand (Franklin et al., 2016), although a few studies have used continuously moving targets. Franklin and Wolpert used smooth hand feedback perturbations in a series of experiments (Franklin & Wolpert, 2008), although they did not focus specifically on the effect of continuous target movement. In a series of studies, Brenner and Smeets investigated reaching to moving virtual targets (Brenner & Smeets, 1994, 1996; Smeets & Brenner, 1995) and reported that response times were slightly higher when targets moved continuously compared to when they jumped instantaneously. Numasawa and colleagues (Numasawa et al., 2022) also investigated the effect of various target movement speeds (not including instantaneous jumps) on reaching corrections and found that, for a range of target speeds, correction latency was unaffected.

Here, we report the results of an experiment that bridges the physical/virtual and moving/jumping target parameters used in previous studies by using physical objects that move in two categorically different ways. In one condition, participants reached toward a physical object that was illuminated; switching the illumination to another, identical object in a different position affected an immediate physical object “jump” similar to common virtual target paradigms; in a second condition, the object moved continuously by rotating about a high-speed motor (similar to that used in Pruszynski et al. 2016 and Reschechtko & Pruszynski 2020). In contrast to previous findings using virtual targets, we found that participants updated their reach trajectories more quickly when reaching toward objects that moved continuously compared to those that jumped instantaneously.

## Materials and Methods

### Participants

20 individuals (ages 19-39; 10 men and 10 women) participated in these experiments. Three participants were left-handed and the rest were right-handed based on self-report; participants used their dominant hand for all experimental procedures. All participants reported normal or corrected to normal vision and no injuries or neurological conditions that would prevent them from making rapid reaching movements. All participants provided informed consent in accordance with procedures approved by the Health Sciences Research Ethics Board at Western University.

### Procedures

Participants sat at a table facing the experimental apparatus. During each experimental trial, the participant used their dominant hand to reach from a stationary starting position to a physical target. The target was a table tennis ball, which was illuminated from within with a white LED. Participants began each trial with the index finger of their dominant hand pressing a switch; when the participant lifted their finger to reach, they released the switch, which initiated target movement following a 30 ms fixed delay. Before initiating the reaching movement, participants received an auditory cue indicating that they could begin reaching toward the initial target at any time. Participants were instructed to reach as quickly as possible and received a second auditory cue 300 ms after they initiated their reach for pacing.

For continuous object movements, the object (i.e. the illuminated table tennis ball) was mounted on the end of a carbon fiber rod which was rotated at high speed from its initial position to final position using a stepper motor with closed-loop control (34HE46-6004D-E1000, OMC Corporation Ltd, China), similar to a previous apparatus described in Pruszynski et al. (2016) and Reschechtko & Pruszynski (2020). The object always started in a central position, from which it could move left (counterclockwise rotation, CCW), right (clockwise rotation; CW), or not move (NM) during catch trials. Movement onset was triggered by the microcontroller 30 ms after the participants lifted their finger from the starting position; in this condition, the motor rotated 15 degrees in 50 ms. The position of each participant’s head was not controlled specifically, but this procedure resulted in the target moving across a visual angle of ~12°, and therefore an average visual velocity of around 240°/sec.

For jumping target movements, LED illumination switched from one stationary object to another object target. These objects were positioned along an arc with radius of 35 cm, directly behind potential target positions for the continuous movement condition. We used this arrangement so that the objects for each stimulus condition provided a very similar visual impression. The arrangement of objects on the apparatus is shown in Fig 1A. When participants triggered target movement for this condition, illumination for the initial (central) object would turn off and illumination of one of the flanking objects (left; CCW, or right; CW) would turn on with the same 30 ms latency as in the moving target condition. In the equivalent of no movement trials, the illumination did not switch.

**Figure 1:**
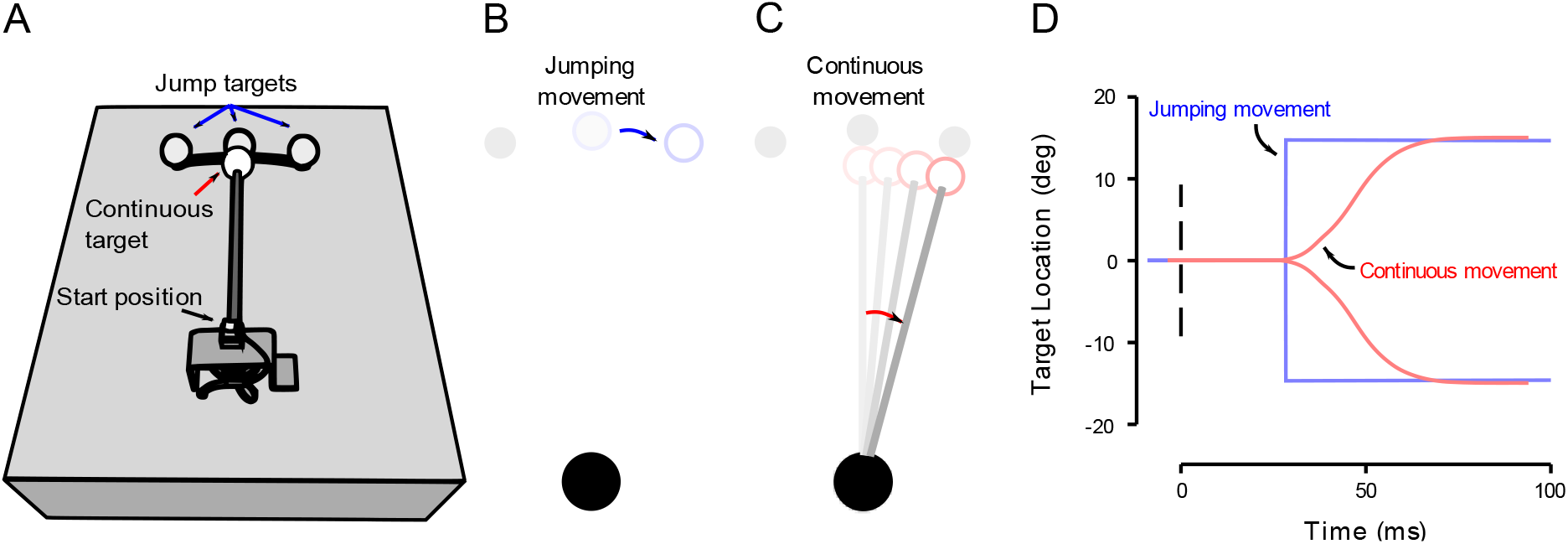
Experimental apparatus and target kinematics. *A:* drawing of the experimental apparatus. *B:* schematic of jumping target movement. During jumping target movements, the central “jumping” target was illuminated from within; on some trials, after participants initiated a reach, the central target turned off and a different target illuminated simultaneously (rightward or “CW” target jump illustrated). The moving object (on the rod) was moved out of the workspace during jumping target trials and did not move during these trials. *C:* schematic of continuous object movement trials. The object on the rod (“continuous target”) began in a central initial position; during some trials, the rod rotated to move the target to a new position directly in front of one of the other jumping target positions (“CW” rotation illustrated). During continuous movement trials, the moving object was illuminated from within and the jumping targets were not illuminated. *D:* schematic of target kinematics. In the continuous movement target paradigm, the object moved to ±15° eccentricity over 50 ms (red traces); in the jumping movement paradigm, the target moved immediately (blue traces). In both paradigms, target movement began 30 ms after participants began their reach (t = 0).

Participants performed 180 reaches to the continuously moving object and an additional 180 reaches to the jumping target. The target conditions interleaved into 60-trial blocks; within each block, participants experienced 20 CW, 20 CCW, and 20 NM trials in a completely randomized order. Half of the participants began with the continuous movement condition (i.e. 60 continuous, 60 discrete, 60 continuous, 60 discrete, 60 continuous, 60 discrete) and the other half began with the jumping movement condition.

### Kinematics

We recorded upper limb kinematics at 120 Hz using optical motion capture (OptiTrack V120:Trio; NatrualPoint Inc, Corvallis, OR). We attached retro-reflective markers to each participant’s dominant arm on the following landmarks: fingernail of the index finger (fingertip); metacarpal-phalangeal joint (MCP) of the index finger; ulnar styloid (wrist); lateral epicondyle (elbow), and acromion process (shoulder); we identified these landmarks using visual inspection and palpation. Primary analyses were carried out using the position of the marker on the index finger (in agreement with previous studies), but some exploratory analyses involve measures related to the position of other arm segments.

### Analysis

To determine the time when participants began to correct their reaching movements (time of divergence), we constructed a time series of receiver-operator characteristics (ROC) for fingertip velocity in each participant and condition. We then determined the onset of corrective movements by finding when the ROC reached certain threshold values. We define the time of divergence as the last time the ROC crossed 60% discriminability (i.e. correctly differentiating between 60% of left- and rightward movements) before reaching a threshold of correctly differentiating between 75% of movements to left- and rightward targets. We considered a variety of ROC criteria for determining divergence time and they yielded similar results.

To investigate potential differences in the gain of movement corrections, we analyzed fingertip position for each participant from 50-100 ms following the time their movements were determined to diverge via ROC. We compared the difference in fingertip position between reaches following leftward target movements to those following rightward target movements to see if this distance was affected by the type of target movement.

We carried out paired t-tests to compare differences in behavior during the discrete and continuous target movements, using the *ttest_rel* function in Scipy library (Virtanen et al., 2020) in Python (Python Software Foundation, Beaverton, OR.

## Results

Participants successfully completed trials without notable exception. Completion of the entire experiment (360 reaches) usually took around 45 minutes. Although the reaches were unconstrained, most fingertip movement was limited to the task-relevant (transverse) plane.

### Participants began to correct movements earlier when the target moved continuously

The primary question we set out to address was whether reach corrections toward continuously moving objects differed from corrections toward targets that move instantaneously. The latency when the object moved continuously (median = 125 ms) was lower (faster) compared to instantaneous target movement (median = 146 ms). These times are generally similar to those observed and reported in previous literature using jumping target movement paradigms (Brenner & Smeets, 1997; Day & Lyon, 2000; Franklin & Wolpert, 2008; Prablanc & Martin, 1992; Smeets & Brenner, 2003) and are also similar to those we reported previously using a physical object (Reschechtko & Pruszynski, 2020). These results are summarized in terms of the heading of the velocity vector in Fig 2A; traces diverge earlier for continuous movement traces. A paired t-test on these divergence times confirmed that reaches toward continuously moving objects consistently diverged earlier (T_19_ = 3.16; P = 5.15e^-3^). Overall divergence times for the two conditions are shown in Fig 2B.

**Figure 2:**
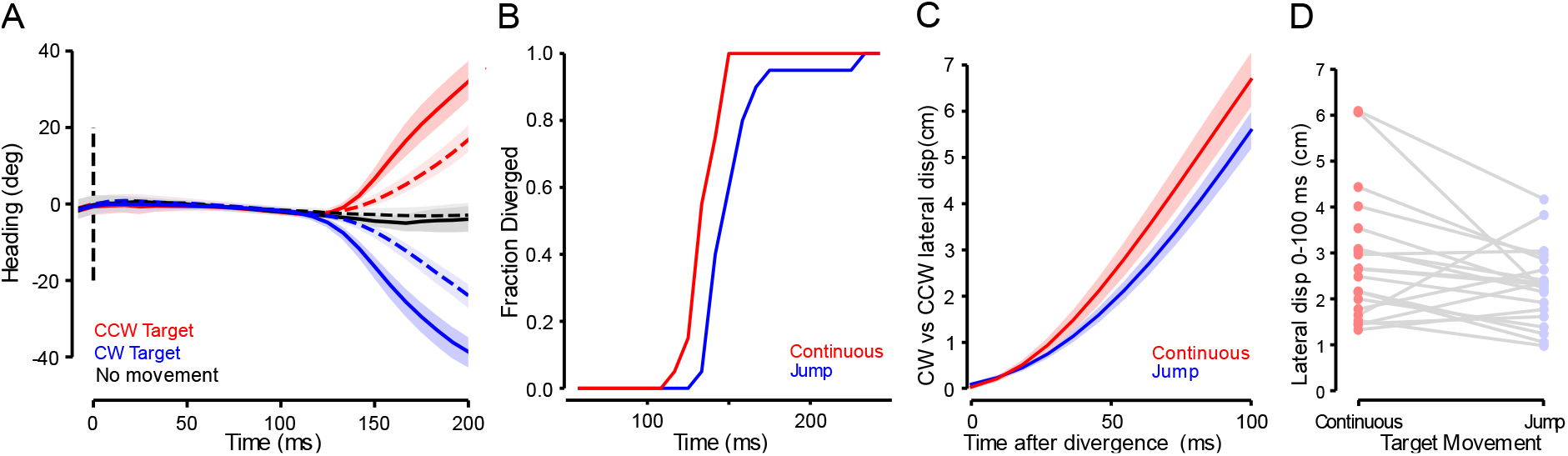
Kinematics, divergence times, and positional differences for reaches toward targets that move continuously or jump. A: main kinematic results. Across-participant average heading angles toward targets that changed position (red: leftward target movement; blue: rightward target movement) and trials where the target did not move (black) for jumping movements (dashed lines) and continuously moving objects (solid lines). Traces are aligned by onset of the target movement; shaded regions are SEM (N=20). *B:* Cumulative distribution function of divergence time when for participants (N = 20) reaching toward an object that moves continuously to a new position (Red) or reaching toward a matched target that “jumps” discontinuously to that position. Divergence time is when reaching trajectories toward leftward and rightward target jumps can be correctly discriminated 75% of the time. *C:* across-participant (N = 20) average differences between fingertip position between movements to rightward and leftward reach corrections toward continuous (red) and jumping (blue) movements. These data are aligned by the onset of divergence for each participant, as determined using the ROC method described in methods. Shaded regions represent SEM. *D:* Paired comparisons (N = 20) of average differences between leftward and rightward reach trajectories during continuous and jumping movements.

**Figure 3:**
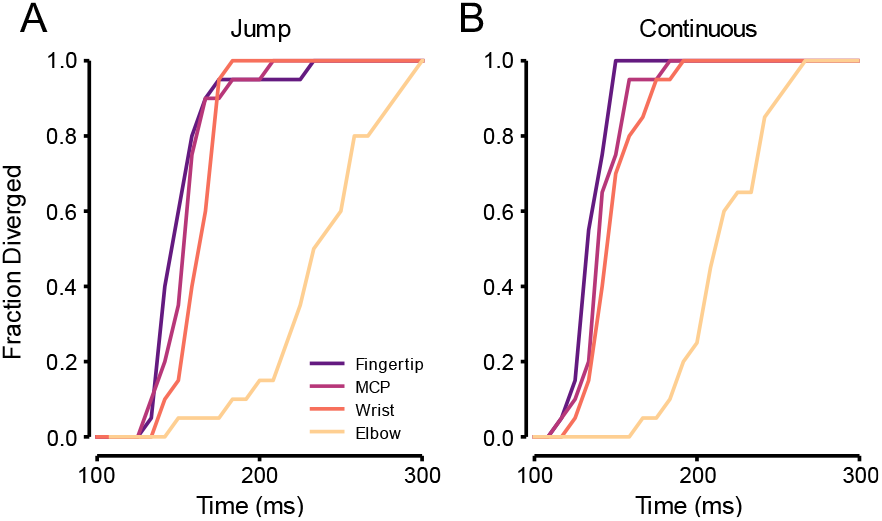
divergence times for all arm segments. *A:* cumulative distribution functions for divergence times computed from ROCs for Jumping target movements for the fingertip (as in fig 2), metacarpal phalangeal (MCP) joint of the index finger, wrist, and elbow of the reaching arm. *B:* divergence times for arm segments in the Continuous target movement paradigm.

### Visual gain was slightly higher for continuous target movement

After determining that people began to change their reach trajectories faster for continuous movements, we investigated whether the magnitude of correction after the time of divergence was different depending on movement paradigm. For each participant, we computed the average position of their fingertip in leftward versus rightward reaches for the 100 ms following the time at which their reaches diverged. On average, the difference in displacement between leftward and rightward reaches was larger when participants were reaching toward the continuously moving target (e.g. higher gain). The average gain in fingertip position from 50-100 ms following time of divergence was higher for the continuously moving target than the jumping target (4.75 ± 0.49 cm compared to 3.81 ± 0.30 cm; paired t-test: T_19_ = 2.12; P = 0.047). This means that, even accounting for the earlier onset of the correction toward a continuously moving object, participants moved their fingers more rapidly when correcting toward a continuously moving object. Fig 2C shows the temporal evolution of difference in fingertip position between movements to left- and rightward target positions aligned by time of divergence, while Fig 2D shows participants’ average differences over 1 00 ms following the time of divergence.

### Relative onset of divergence is uniform across upper limb segments

Due to the kinematic redundancy in this unconstrained task, it was possible for people to make corrective actions without all segments of the upper limb moving in a correction-specific direction. We thought that people might change their coordination strategies in the different movement paradigms because we previously observed some participants who appeared to make a final correction with the hand only, while the wrist appeared to remain relatively stationary via wrist ab/adduction. Therefore, we tracked the positions of various parts of the arm during this study and examined whether the corrective action is implemented by a proximal arm segment (e.g. movement about the shoulder), or a distal one (e.g. movement about the elbow or wrist). One potential reason people might select one strategy versus another is that the distal segments have lower inertia; on the other hand, descending commands reach proximal musculature more quickly than distal musculature.

We carried out an exploratory analysis by computing divergence times (as we did for the fingertip) for markers attached to the MCP, wrist, and elbow joints. Using ROC curves, it was possible to distinguish between movements toward leftward and rightward target movements from all segments and for both movement paradigms. In general, divergence times were later for segments that were more proximal. For reaches toward jumping targets, median divergence times were 142, 150, 158, and 229 ms following target movement onset for the fingertip, MCP, wrist, and elbow markers, respectively. For reaches toward continuously moving objects, median divergence times were 125, 133, 142, and 208 ms following target movement. Overall, there was no clear effect of the target movement paradigm on arm segment use: we observed the same relationship (more rapid divergence toward objects that moved continuously than targets that jumped) regardless of the segment we analyzed.

Finally, we investigated whether, within individual participants, earlier divergence at one segment was related to earlier divergence at another segment by correlating divergence times across each pair of arm segments for each target condition. Divergence times at the fingertip and MCP joint were positively correlated for reaches to jumping targets (*r* = 0.87; *P* = 4.93e^-7^) and continuously moving objects (*r* = 0.59; *P* = 0.006), and divergence times for the MCP and wrist were positively correlated during reaches to jumping targets (*r* = 0.5; *P* = 0.024), but none of the other correlations were significant. Across target movement paradigms, elbow divergence was usually the latest of all segments (all participants for the moving object and 18 for the jumping target). While the order of divergence recapitulated the median times of divergence (most participants diverged at fingertip, then MCP, and then wrist), there was notable variability. When reaching toward the moving target, four participants showed earliest time of divergence at the wrist for the moving object paradigm; however, when moving toward the jumping target, only one of those participants showed earliest divergence at wrist – even though four participants overall showed earliest time of divergence in the wrist for this movement paradigm as well. These results likely point to the kinematic redundancy that people naturally exploit during this unconstrained task, despite its rapid timescale.

## Discussion

Our results indicate that, at least under certain circumstances, people correct ongoing reaching movements more quickly if the object they are tracking shifts continuously compared to when it jumps. This may occur because different target movement paradigms induce different eye movements to track target movement. Curiously, these results are not consistent with previous results employing virtual targets (Brenner & Smeets, 1996; Numasawa et al., 2022; Smeets & Brenner, 1995, 2003), which potentially indicates the interaction of target movement paradigms with the realism of the target.

### Eye movements and arm movements

When the eyes fixate on an object and that object begins to move, the movement of the object on the retina (“retinal slip”) drives smooth pursuit eye movements to bring the object back toward the fovea smoothly (Krauzlis, 2004; Ono, 2015). In contrast, when a target jumps to a new position instantaneously, the target shift drives saccadic eye movement in which the fovea is shifted ballistically to the new position (Leigh & Zee, 1999). Although we did not record eye movement in this experiment, we expect that the continuous object movement paradigm we used elicited pursuit eye movements, whereas the jumping paradigm elicited saccades.

Onset of smooth pursuit movements is reportedly 25-75 ms earlier than that of saccades (Adler et al., 2002; Krauzlis & Miles, 1996). A number of reasons for this observation have been proposed, including a speed-accuracy tradeoff in which pursuit is initiated more quickly but is less accurate; the increased salience of continuously moving targets (although the basis for this contention is unclear); and increased complexity of saccade preparation due to additional spatial transformations being required (Sparks & Mays, 1990). Rapid reaching corrections likely involve subcortical circuits (Alstermark et al., 1987; Day & Brown, 2001), and those circuits could be related to some of those involved in the production of smooth pursuit. If some of the same circuits are involved in reach corrections and eye movement, the increased speed of reach corrections to continuously moving targets could be a product of the same processes that yield faster responses during pursuit eye movement. The moderate increase in correction gain we observed could also be evidence for increased salience of continuously moving targets: overall, corrections toward the continuously moving objects both started earlier and yielded more rapid arm movement (Fig. 2).

Another potential reason for differences in reach speeds is related to availability of visual information. In our study’s continuous movement condition, information related to both position and motion were available, whereas only position information was available in the jumping target condition. A body of visuomotor research has investigated the contributions of feedback on position and motion on reach corrections (Blouin et al., 1993; Paillard, 1996; Saunders & Knill, 2003, 2004). These studies, inspired by contributions of magno- and parvocellular visual pathways through the lateral geniculate nucleus (Lennie, 1980) have focused on the interaction between visual target motion and location in the visual field and latency of correction. In contrast to the simple idea that additional information (e.g. regarding target motion) would increase correction speed, a common proposal from these studies is that the peripheral visual field is particularly sensitive to rapid motion used for early reach corrections, whereas the central visual field (<15° eccentricity) is specialized for position information to aid in arriving at a final position (but see Saunders & Knill, 2004). Our experimental paradigm does not neatly fit into this literature, both because we delivered rapid stimulus movement to the central visual field, and because our targets stopped moving early in the reach, rather than persisting throughout the reach.

### Physical and virtual, moving and jumping targets

To some extent, the present results are opposed to those reported in previous work comparing reaching movements to virtual targets that move instantaneously or continuously. Brenner and Smeets conducted a series of studies (Brenner & Smeets, 1997, 1996; Smeets & Brenner, 2003) that investigated the effect of several aspects of target movement on reaching behavior. Among the factors investigated were a comparison of movement speed, including contrasting jumping and continuously moving targets. They found that people made somewhat faster corrections when targets moved faster — and corrections were more rapid when the targets jumped compared to when they moved continuously. There are a variety of differences between our experimental paradigm and those previously mentioned. Brenner and Smeets used targets which moved continuously from the time they appeared (rather than beginning in a stationary position and then beginning to move), and they instructed their participants to intercept those targets during movement. In contrast, our continuously moving object began stationary and moved more rapidly such that it was finished moving by the time participants had initiated their reach corrections. In addition, participants in Brenner and Smeets’ experiments used tools (pointers, computer mice) and reached toward virtual targets; as such, the tools they used were passive implements that could not grasp physical targets, and the targets themselves were not graspable. Graspability (as determined by effector and/or target) could be important for modulating the neural circuits involved in the action (Freud et al., 2018; Gomez et al., 2018); additionally, it is possible that the additional complexity of anticipating the target velocity for interception in renner and meets’ paradigm delayed reaching corrections.

Numasawa et al. (2022) also investigated the effect of different target moving speeds on reaching corrections. Although they did not compare the effect to a jumping target, they did not observe differences in correction latency depending on target velocity. In contrast, they report that gain was higher for higher target velocities. In comparison to the current paradigm, however, Numasawa and colleagues used target velocities that were much slower: 10°/s-40°/s compared to the average of 240°/s we used, and the onset latencies they observed were around 50 ms longer than ours. These longer onset times are similar to those observed when people integrate additional instructions into reach corrections (reaching in the opposite direction: Day & Lyon 2000; Reschechtko & Pruszynski 2020; stopping an ongoing reach: Pisella et al. 2000), which, compared to rapid reaching corrections, are likely to engage higher-order neural circuits at the cost of response time. If these movements did not rely on subcortical circuitry specific to rapid reach corrections, any overlap between these circuits and those underlying pursuit eye movements may have been irrelevant.

As previously mentioned, the discrepancy between the effect of jumping and moving targets in our paradigm compared to previous results could related to our use of physical targets rather than the virtual targets used in previous studies. Physical objects elicit different neural activity than virtual objects even when these objects are merely presented to the field of view (Fairchild et al., 2021; Gallivan et al., 2009). Further, placement of physical objects within the workspace, or “graspable” objects, (Gomez et al., 2018) appears to additionally modulate brain activity, suggesting that responses we see in the present study could take advantage of this specialization of the visuomotor system. The motor system is presumably specialized to interact with physical objects as we reach for them, so it is possible that physical objects, moving in physically possible ways, are more salient than other movement patterns. Both the lower latency and higher gain we observe in movements toward continuously moving objects could be evidence for increased salience. The naturalism in our paradigm is also increased compared to many previous studies because people are using their hand to point at a physical and reachable object, rather than moving a tool to point toward an unreachable virtual target.

A primary consideration in the study of natural movement should be the use of relatively natural stimuli. While computer and projection screens allow for the convenient delivery of target stimuli, evidence continues to build that neural responses can differ based on stimulus properties including their reachability and realism (Snow & Culham, 2021). Our results, when contrasted with the existing literature on reaching corrections to virtual targets, provide evidence that neural responses underlying reaching corrections are also different depending on the use of virtual targets or physical objects. Development of apparatus to better control the delivery of virtual and physical targets in more similar ways in the same group of participants could further test the robustness of this comparison.

## Limitations

A number of considerations make it difficult to directly compare different target kinematics across physical objects and virtual targets. One primary difficulty is matching enough parameters that the physical object and virtual target paradigms are reasonably similar. The virtual target paradigms we have cited here involve different reach distances, instructions, stimuli, and effectors. These differences make it difficult to compare our results to previous results, and to know whether our results generalize to a wider range of movement velocities, target sizes, and so forth. A second consideration is that we did not record eye movement, so connections between arm and eye movement are based only on the differences in movement paradigm and eye movements similar paradigms have evoked. Although the onset of hand movement often follows the onset of eye movement (Georgopoulos et al., 1981; Land & Hayhoe, 2001; Neggers & Bekkering, 2000), there are cases when hand movement can precede eye movement (Abekawa et al., 2014); nonetheless, even in those cases, onset of eye and hand movements are highly correlated, suggesting that time differentials in eye movement onset in smooth pursuit and saccades are likely to be conserved in reaches as well. Finally, the physical movement of the object during continuous movement produced some audible noise when the motor moved the target; while this noise is not direction-specific, we cannot rule out the possibility that it affected response latency.

## Acknowledgements

This work was funded by a Canadian Institutes of Health Research (CIHR) Foundation grant (to J.A.P.). The Optitrack camera system was purchased with a CIHR research stipend to S.R. C.G. received a salary award from the Natural Sciences and Engineering Research Council (NSERC USRI). S.R. received salary awards from CIHR (postdoctoral fellowship) and Western BrainsCAN through the Canada First Research Excellence Fund (CFREF). J.A.P. received a salary award from the Canadian Research Chair program.

## Disclosures

No conflicts of interest, financial or otherwise are declared by the authors

## Author Contributions

S.R. and J.A.P. conceived and designed research, C.G. and S.R. performed experiments; C.G. and S.R. analyzed data; S.R. and J.A.P. interpreted results of experiments; S.R. prepared figures; S.R. drafted manuscript; S.R. C.G. and J.A.P. edited and revised manuscript.

## Notes

### Competing Interest Statement

The authors have declared no competing interest.

### Summary of Updates

-Changed language to describe moving "objects" and jumping "targets" -Revised citations errors -Added discussion paragraph regarding motion and position information (page 10, first full paragraph).

